# Chronic exposure of mice to phthalates enhances TGF beta signaling and promotes uterine fibrosis

**DOI:** 10.1101/2023.05.10.540240

**Authors:** Ritwik Shukla, Arshee Mahmuda, Mary J. Laws, Jodi A. Flaws, Milan K. Bagchi, Amy J. Wagoner Johnson, Indrani C. Bagchi

**Author notes:** Contact: To whom correspondence should be addressed: Indrani C. Bagchi. Disclosure Statement: The authors have nothing to disclose.

## Abstract

Phthalates are synthetic chemicals widely used as plasticizers and stabilizers in various consumer products. Because of the extensive production and use of phthalates, humans are exposed to these chemicals daily. While most studies focus on a single phthalate, humans are exposed to a mixture of phthalates on a regular basis. The impact of continuous exposure to phthalate mixture on uterus is largely unknown. Thus, we conducted studies in which adult female mice were exposed for 6 months to 0.15 ppm and 1.5 ppm of a mixture of phthalates containing di(2-ethylhexyl) phthalate, di-iso-nonyl phthalate, benzyl butyl phthalate, di-n-butyl phthalate, diisobutyl phthalate, and diethyl phthalate via chow ad libitum. Our studies revealed that consumption of phthalate mixture at 0.15 ppm and 1.5 ppm for 6 months led to a significant increase in the thickness of the myometrial layer compared to control. Further investigation employing RNA-sequencing revealed an elevated transforming growth factor beta (TGF-β) signaling in the uteri of mice fed with phthalate mixture. TGF-β signaling is associated with the development of fibrosis, a consequence of excessive accumulation of extracellular matrix components, such as collagen fibers in a tissue. Consistent with this observation, we found a higher incidence of collagen deposition in uteri of mice exposed to phthalate mixture compared to unexposed controls. Second Harmonic Generation imaging showed disorganized collagen fibers and an increase in uterine stiffness upon exposure to phthalate mixture. Collectively, our results demonstrate that chronic exposure to phthalate mixture can have adverse effects on uterine homeostasis.

## 1. INTRODUCTION

Phthalates, esters of phthalic acid, are synthetic chemicals widely used as plasticizers in various consumer products (1–5). The high molecular weight phthalates with a long-chain include di(2-ethylhexyl) phthalate (DEHP), di-iso-nonyl phthalate (DiNP), di-iso-decyl phthalate (DiDP), di-n-octyl phthalate (DnOP), and di(2-propylheptyl) phthalate (DPhP). They are used as a part of polyvinyl chloride. Phthalates with a short chain are used in the manufacture of personal care products, such as perfumes, nail polish, deodorants, and lotions. Phthalates that belong to this category include dimethyl phthalate (DMP), diethyl phthalate (DEP), benzylbutyl phthalate (BBzP), di-n-butyl phthalate (DnBP) and di-iso-butyl phthalate (DiBP).

An enormous amount of phthalates are used worldwide each year (6). Since phthalates readily leach into the environment from plastic products because they are not covalently bound to the plastic, humans are constantly exposed to various phthalates through inhalation, ingestion, and dermal contact (7). Ingestion is the most common route of exposure (8,9). Studies indicate that phthalate metabolites are present in nearly 100% of tested human urine samples (10–12). Interestingly, phthalate metabolite levels are higher in women than men. This is likely due to higher use of personal care products by women compared to men (13).

Increasing scientific evidence links phthalate exposure to harmful health outcomes due to endocrine-disrupting properties displayed by this class of chemicals. Epidemiological studies indicate that exposure to phthalates is associated with decreased pregnancy rates, high miscarriage rates, and preeclampsia in women (10,14–19). Phthalate exposures are also associated with adverse pregnancy outcomes including low birth weight and impaired childhood intellectual development and growth (15,17). The mechanism by which phthalate exposure impacts uterine function and causes pregnancy loss in women is not known.

Studies using rodent models provide insights into the physiological processes by which phthalates interfere with reproductive functions. Previous reports indicated that prenatal exposure to phthalates affects folliculogenesis, delays puberty and decreases fertility (20,21). It was also reported that exposure to DEHP causes uterine abnormalities in mice, such as decreased luminal epithelium proliferation and increased numbers of abnormally dilated blood vessels in the endometrium (22). Our previous study showed that exposing pregnant mice to an environmentally relevant level of DEHP did not impact pregnancy unless the mice were fed a high fat diet (23). A recent study showed that exposure to DiNP, at an environmentally relevant dose compromises the ability of mice to maintain pregnancy; about 33% of the females exposed to this dose of DiNP were unable to maintain pregnancy (24,25). While these studies have focused on exposure to a single phthalate over a limited period, humans are exposed to a mixture of phthalates daily over a lifetime.

Thus, we conducted a study in which mice were exposed long-term to an environmentally relevant level of phthalate mixture via chow ad libitum and analyzed the effects of this exposure on the uterus. The phthalate mixture composed of DEHP, DiNP, BzBP, DBP, DiBP, and DEP, was designed based on the Illinois Kids Development Study (I-KIDS) in which levels of phthalate metabolites were measured in the urine samples of pregnant women (26). A recent study reported that chronic dietary exposure to this phthalate mixture alters estrous cyclicity (27). However, the effects on the uterus remain unclear. In this study, we tested the hypothesis that long-term exposure to the phthalate mixture alters uterine homeostasis. We performed molecular analysis, imaging, and measured tissue stiffness to gain insights into the impact of phthalates on uterus.

## 2. Materials and methods

### 2.1. Chemicals

The phthalates were purchased from Sigma-Aldrich (St. Louis, Missouri) and had 98% purity. Corn oil (Columbus Vegetable Oils, Des Plaines, Illinois) was used as the vehicle control. The phthalate mixture was prepared and diluted in tocopherol-stripped corn oil (vehicle control).

### 2.2. Animals

Female CD-1 mice at 33 days of age and male CD-1 mice at 7 weeks of age were purchased from Charles River Laboratories (Wilmington, Massachusetts) and housed in the College of Veterinary Medicine vivarium at the University of Illinois Urbana-Champaign (Urbana, Illinois). As previously described, female mice were group housed 3 mice per cage in polysulfone cages (Allentown, Allentown, New Jersey), with 1/8 corn cob bedding (Shepherd Specialty Papers), environmental enrichment (iso-BLOX, catalog #6060, Envigo), and water purified by reverse osmosis (27). The University of Illinois Institutional Animal Care and Use Committee approved all animal handling, housing, and procedures.

### 2.3. Study design and dosing

Phthalates were administered to mice *ad libitum* in the rodent chow as previously described (27). The base chow was a modification of the AIN-93G formulation (TD.94045) that replaced soybean oil with corn oil (Columbus Vegetable Oils, Des Plaines, Illinois). Chow with 7% corn oil was used as the vehicle control group. Phthalates were mixed in corn oil and provided to Envigo for chow preparation. The phthalate mixture used in this study consisted of 35 % DEP, 21 % DEHP, 15 % DiNP, 15 % DBP, 8% DiBP, and 5% BzBP. This environmentally relevant phthalate mixture was developed based on phthalate metabolite concentrations in urine of pregnant women in central Illinois (26). High levels of phthalates are found in food contact materials, such as plastic wrap and plastic containers. Thus, ingestion is a major route of exposure for humans (9). To mimic human exposure, phthalate mixture exposure was administered via the rodent chow. Phthalate mixture doses used in the study were 0.15 ppm and 1.5 ppm. Each treatment group consisted of 12–14 mice. Based on previous studies that exposed mice to phthalates via the chow, target doses were selected based on the assumption that a 25 g mouse eats approximately 5 g of food/day (28–30). Accordingly, the 0.15 ppm dose is roughly equivalent to 24 mg phthalate/kg body weight/day and the 1.5 ppm dose is roughly equivalent to 240 mg phthalate/kg body weight/day. The 0.15 and 1.5 ppm doses fall within the ranges of daily human exposure, infant exposure, and occupational exposure (31,32). Beginning at 6 weeks of age, the mice were administered treatments in the chow *ad libitum*, continuously for 6 months. Daily continuous exposure to phthalates strongly mimics human exposure to phthalates.

### 2.4. Tissue collection and evaluation of uterine histology

At 6 months of age, mice were euthanized in diestrus, and uterine horns were collected. A part of the uterine horn was fixed in 10% neutral buffered formalin (NBF) fixative while the other part was flash frozen in liquid nitrogen for gene expression analysis. For histological analysis, fixed uteri were dehydrated using a series of increasing concentrations (70% to 100%) of ethanol. Samples were embedded in paraffin, sectioned, and stained with hematoxylin and eosin. The thickness of the myometrial layer in the uterine sections were determined by random measurement of myometrial layer at 8 random locations.

### 2.5. RNA sequencing and gene expression analysis

Frozen uteri collected at 6 months from control and 0.15 ppm phthalate mixture treated groups were used for RNA sequencing. Total RNA was extracted from uterine horns using a standard TRIzol-based protocol. RNA integrity was verified using Agilent 2100 bioanalyzer (Agilent Technologies Inc., Santa Clara, CA, USA) and sequencing was performed at the Biotechnology Center of the University of Illinois, Urbana-Champaign. A list of genes that had a relative fold change of >2 was further sorted by gene ontology and pathway analysis.

For gene expression analysis, RNA was converted to cDNA and real-time quantitative PCR reactions were carried out using SYBR-green master mix (Applied Biosystems) on a QuantStudio^TM^ 3 Real-time PCR instrument (Applied Biosystems). The mean threshold cycle (Ct) for each sample was calculated from Ct values obtained from three replicates. The normalized ΔCt in each sample was calculated as the mean Ct of the target gene subtracted by the mean Ct of the reference gene. ΔΔCt was then calculated as the difference between the ΔCt values of the control and mutant samples. The fold change of gene expression in each sample relative to control was generated using the 2^−ΔΔCt^ mathematical model for relative quantification of quantitative PCR. The mean fold induction and standard error of the mean (SEM) were calculated from at least three independent experiments.

### 2.6. Masson’s Trichrome and Picrosirius red stain

Collagen deposition in the uterine sections was analyzed by staining with Mason trichrome stain and picrosirius red. For Masson’s trichrome staining, 5-micrometer uterine sections were prepared, and the paraffin was removed. The sections were rehydrated using a series of alcohol solutions with varying concentrations, and then washed with distilled water. To improve the quality of staining, the sections were refixed with Bouin’s solution for one hour. To remove any yellow color, the sections were washed under running tap water for 5 to 10 minutes. Next, the sections were stained with Weigert’s iron hematoxylin solution for 10 minutes and rinsed with warm tap water for 10 minutes, followed by a rinse with distilled water. The sections were then treated with Biebrich scarlet-acid fuchsin solution for 10-15 minutes, followed by another rinse with distilled water. They were then placed in a solution of phosphomolybdic and phosphotungstic acid for 10-15 minutes. After that, the sections were transferred to an aniline blue solution for 5-10 minutes, briefly rinsed with distilled water, and then differentiated in a 1% acetic acid solution for 2-5 minutes. Finally, the sections were washed with distilled water, dehydrated quickly using 95% ethyl alcohol, cleared in xylene, and mounted with resinous mounting medium.

The Picrosirius red staining was used to detect the total collagen as previously described (33). The fixed tissues were processed using an automated tissue processor and embedded in paraffin. Uterine sections of 5 μm were obtained using a sliding microtome and attached to a slide glass. The paraffin-embedded sections were deparaffinized three times in xylene for 5 minutes each and then de-alcoholized in a series of ethanol solutions with decreasing concentrations (100%, 100%, 90%, and 70%) and ultrapure water for 5 minutes each. The slides were immersed in a Picrosirius red staining solution for 1 hour. After staining, the slides were rinsed twice in a 0.5% acetic acid solution. The sections were dehydrated with three changes of 100% ethanol and cleared using three changes of xylene. Finally, the sections were mounted using a resinous mounting medium.

### 2.7. Second Harmonic Generation (SHG) imaging and image quantification

A confocal microscope (Zeiss LSM 710, Germany) was used to image paraffin embedded uterine sections. A Ti: Sapphire laser producing 100-fs pulses centered at 780 nm wavelength illuminated the samples. A 10x 0.25 NA objective lens collected the SHG signal to construct the 2D images of the whole cross-sections of the tissue. Then with a 40x 1.2 NA objective water immersion lens, volumetric images (141.7*141.7*20 μm) of the myometrium were obtained. The SHG parameters including out-of-plane fiber angle (ϕ), circular variance (CV), and spherical variance (SV) were computed for each sample using the orientation data provided by FT-SHG analysis (34,35). ϕ represents the average angle of the collagen fibers with respect to the x–y imaging plane, where 0° and 90° indicate in-plane and out-of-plane, respectively. CV is the dispersion of the fiber orientations in the imaging plane with a value between 0 (aligned fibers) and 1 (randomly oriented fibers). Similarly, SV describes the dispersion of fiber orientations in 3D ranging from 0 to 0.5. For example, fully aligned fibers in the imaging plane would have a low value for CV; a volumetric stack of imaging planes with fibers of similar alignment in all the imaging planes would have a low SV; and a stack with fibers oriented parallel to the imaging planes of the volumetric stack would have a low ϕ.

### 2.8. Nanoindentation and co-registration

Frozen uterine horns (n=6) were thawed to room temperature for 5 minutes. Two mm length pieces of tissue were cut transversely from the horns, embedded in an optimal cutting temperature compound, and cryosectioned at −20 °C perpendicular to the uterine lumen. The tissues were cryosectioned before nanoindentation to reduce the surface roughness of the samples. A Piuma nanoindenter (Optics11, Amsterdam, Netherlands) acquired the force-indentation data from which the indentation modulus was determined using the model described previously (36). The indentation modulus quantifies the elastic response of a material subjected to the action of a concentrated load in a single point and it represents the local stiffness of the tissue. The spherical probe radius was 99 μm and the cantilever stiffness was 0.50 N/m. The built-in camera of the nanoindenter captured brightfield images of the entire uterine cross-section. To locate the indentation points on the cross-section, another set of brightfield images was captured right before indentation, when the probe was in contact with the sample. The sample was indented every 50 μm along one medial-lateral line. The maximum indentation depth was 5 μm, with a displacement rate of 5 μm/s. The samples were submerged in buffer solution throughout indentation.

The images of indented samples versus the SHG images were co-registered to identify the local mechanical properties of the tissue as described previously (35). After indenting, the samples were placed on a cover glass and buffer solution dispensed using a pipette kept the sample hydrated during imaging. A set of brightfield images of the whole cross-section was acquired with the SHG images in the LSM microscope. The bright-field images from the nanoindenter and the SHG microscope were co-registered semi-manually using the tissue boundaries as fiducial references. Finally, the indentation data were overlaid on the SHG image using the coordinates taken from the two sets of bright-field images.

### 2.9. Statistical Analysis

Statistical analyses for ImageJ and q-PCR analyses were performed as we have done previously (37). Analyses were done using a two-tailed Student’s *t-test*, Mann-Whitney rank sum test (for single comparison), one-way analysis of variance (ANOVA) with a Bonferroni post-test (for multiple comparisons between samples or time points), or two-way ANOVA with a Bonferroni post-test (for multiple comparisons between different samples and time points). In addition, an analysis of equal variances was done on all numerical data to determine whether a parametric or non-parametric hypothesis test was appropriate. Data were expressed as mean ± SEM and were considered statistically significant at p ≤ 0.05. All data were analyzed and plotted using GraphPad Prism 9.4 (GraphPad Software). The OriginPro 9.7 software package (OriginLab Corporation, Northampton, MA) was used for the statistical analysis of SHG imaging and nano-indentation. A one-way ANOVA determined the differences between the means of different regions, followed by a post hoc Tukey analysis which indicated which groups differed from one another. Three significance levels (p<0.05, p<0.01, p<0.001) was used for the analysis.

## 3. RESULTS

### 3.1. Effect of chronic exposure to a mixture of phthalates on uterine histology

To investigate the effects of long-term, dietary phthalate exposure on the uterus, adult mice were fed chow containing vehicle control (corn oil) or a mixture of phthalates at doses 0.15 ppm and 1.5 ppm. The 0.15 ppm and 1.5 ppm doses fall within the ranges of daily human exposure, infant exposure, and occupational exposure (31,32). Representative hematoxylin and eosin (HE) stained uterine sections obtained from control and mice exposed to phthalate mixture for 6 months were analyzed to study changes in uterine histology. While there was no significant difference in uterine endometrium in any treatment groups, an increase in the thickness of the myometrial layer in response to the mixture was observed in the 0.15 ppm and 1.5 ppm groups compared to controls (Fig. 1).

**Figure 1:**
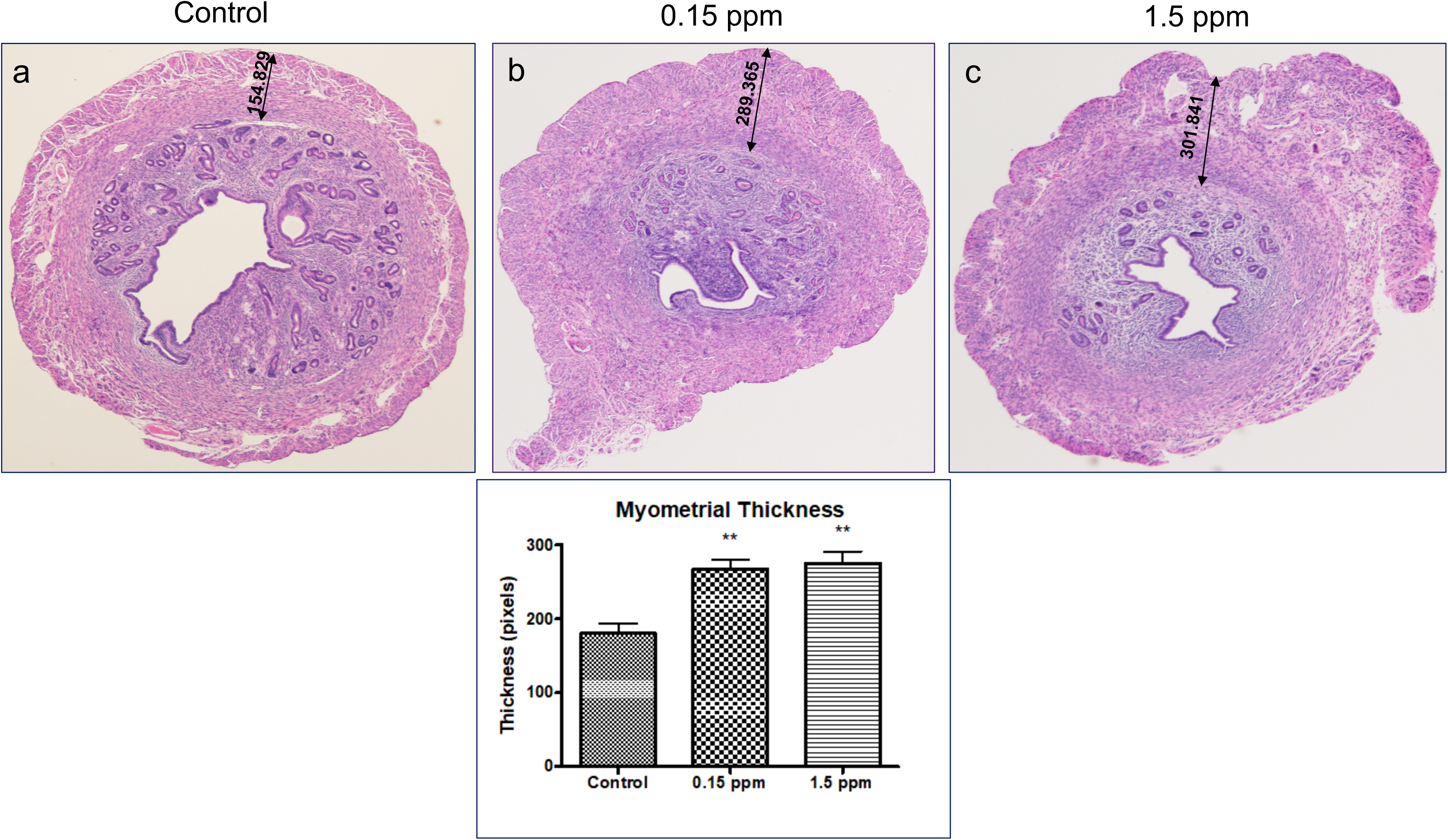
Histological examination of uterine sections. Upper: Serial sections of uterine horns of control (a), 0.15 ppm (b) and 1.5 ppm (c) of phthalate mixture exposed mice at 6 months were examined by H&E staining. The representative images from the control and phthalate mixture treated groups are shown. Lower: Eight random measurements for myometrial layer were determined by ImageJ analysis. Data represent mean ± SEM from three separate samples. Asterisks indicate statistically significant differences (\*\**P < 0*.01).

### 3.2. RNA-sequencing analysis

To obtain insights into biological processes that are impacted upon phthalate exposure in an unbiased manner, we performed RNA-sequencing to compare the expression levels of transcripts in the uteri of control and mice exposed to 0.15 ppm phthalate mixture for 6 months. Interestingly, our study revealed an up-regulation of transcripts involved in the transforming growth factor-beta (TGF-β) signaling pathway in the phthalate-exposed uteri compared to untreated controls (Fig. 2A). The RNA-seq data were validated by real-time PCR analysis. As shown in Fig. 2B, we confirmed a marked up-regulation of mRNA corresponding to TGF-β1, a cytokine that plays a critical role in regulating cell growth, differentiation, and migration in various biological processes, in 0.15 ppm or 1.5 ppm groups (panel a). The expression of biglycan, a proteoglycan that binds to and modulates the activity of TGF-β1, was significantly increased in uterine samples of 0.15 ppm groups (panel b). While the expression of TGF β 1 receptor 1 (TGF-βR1) did not change across samples (panel d), we observed increased expression of integrin beta 1, a transmembrane receptor that can directly interact with the TGF-β receptor and activate downstream signaling pathways, in 0.15 ppm group compared to control (panel c). Collectively, these results indicated that a long-term exposure to phthalate mixture enhances TGF-β signaling in the uterus.

**Figure 2:**
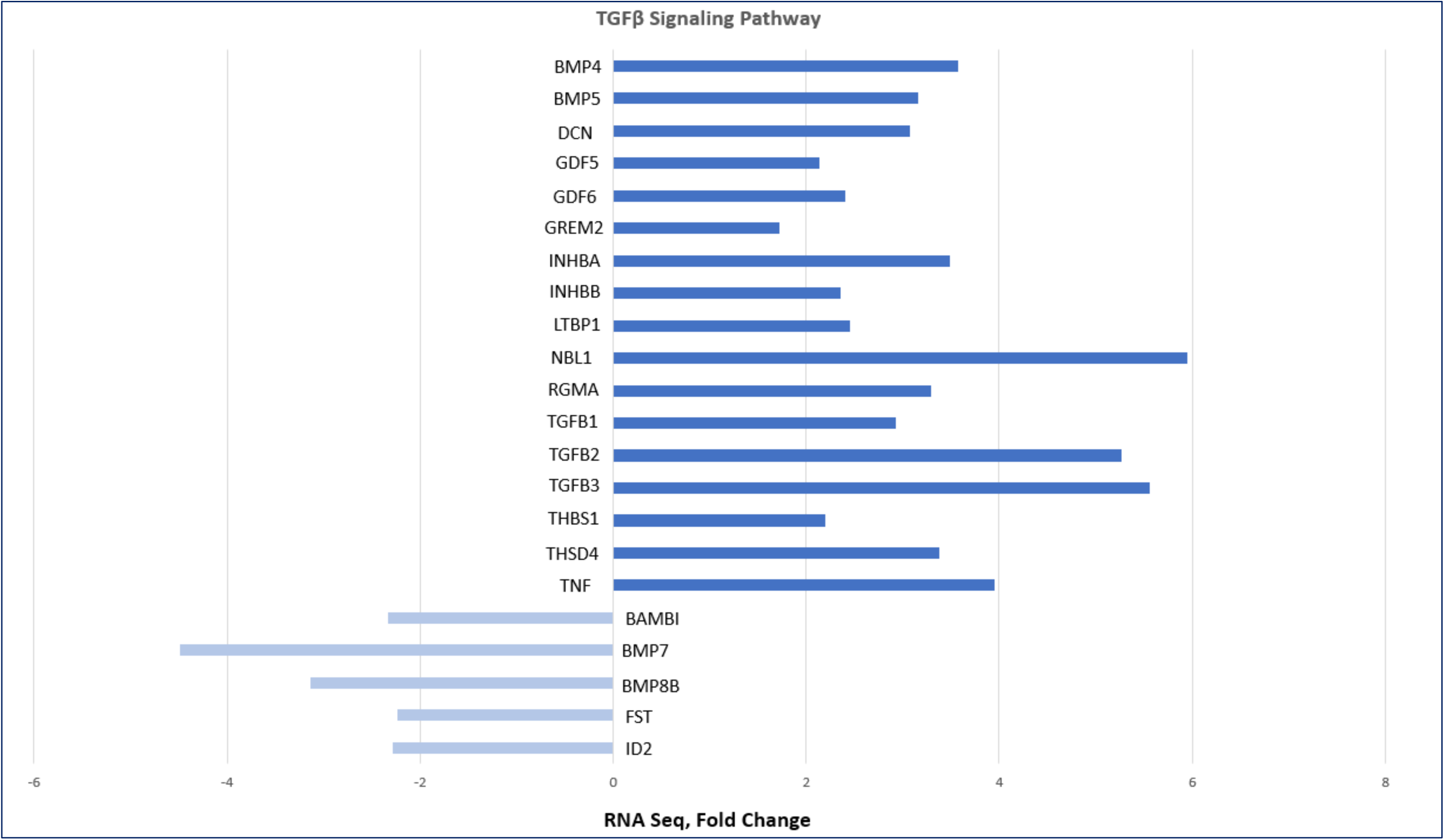
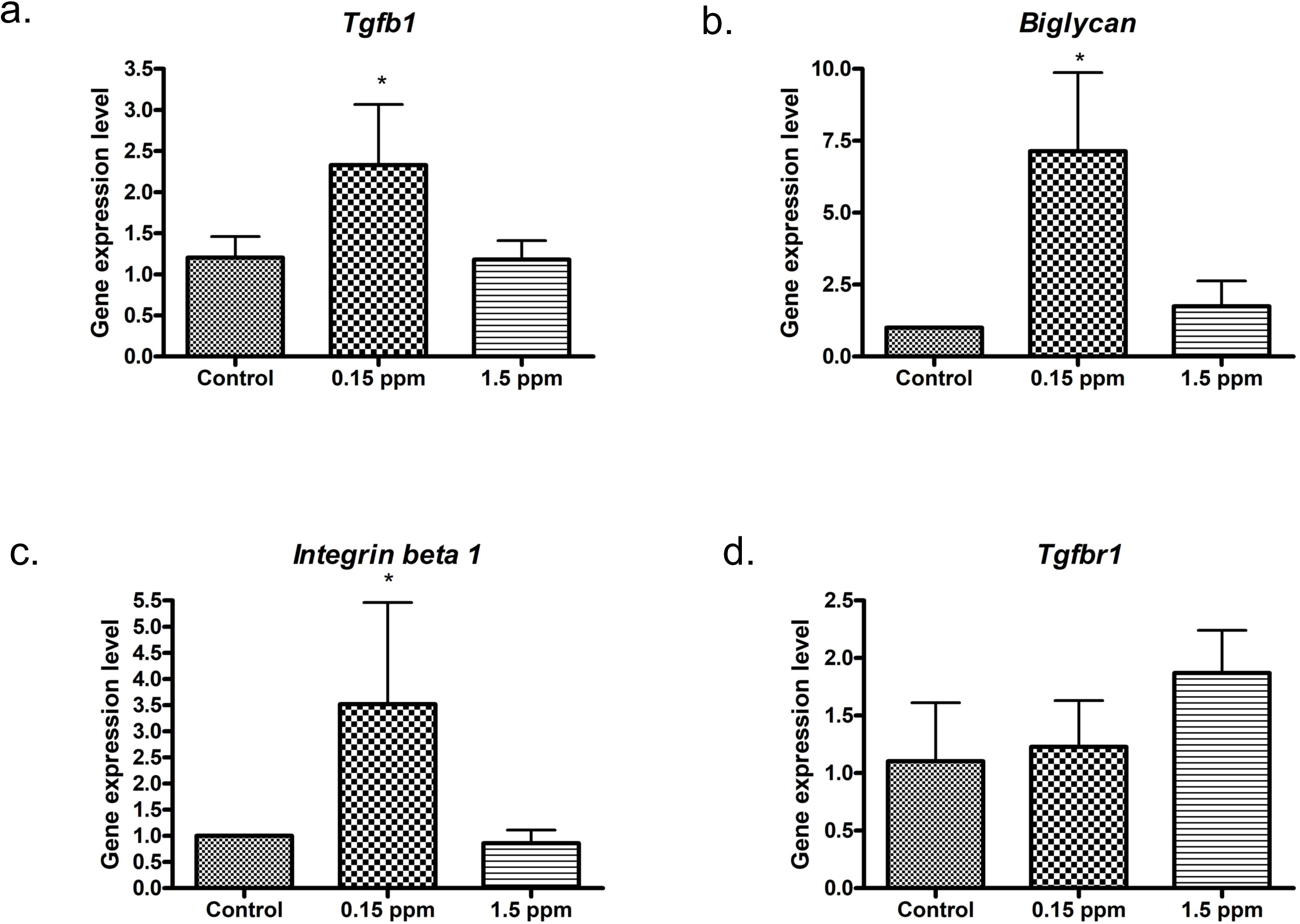
Exposure to a phthalate mixture leads to elevated TGF-β signaling in the uterus. A: Uteri collected at 6 months from control groups and 0.15 ppm groups were used for RNA-seq analysis. Transcriptional changes corresponding to factors in the TGF-β pathway that were identified in RNA-seq of 0.15 ppm groups versus control groups are shown. TGF-β pathway genes that were upregulated in 0.15 ppm groups compared to control groups are shown in dark blue. TGF-β pathway genes that were downregulated in 0.15 ppm groups compared to control groups are shown in light blue. B: Uteri collected at 6 months were used for quantitative real-time polymerase chain reaction (qPCR) analysis. Uterine RNA from control groups, 0.15 ppm groups, and 1.5 ppm groups were subjected to qPCR using primers specific for *Tgfb1* (a), *Biglycan* (b), *Integrin beta* 1(c), and *Tgfb receptor 1* (d). *36b4* was used as the internal control. Data represent mean ± SEM from six separate samples. Asterisks indicate statistically significant differences (\**P < 0*.05)

### 3.3. Expression of collagen family members in the uterus

Elevated TGF-β signaling is associated with the development of uterine fibrosis, a consequence of excessive accumulation of extracellular matrix components, including collagen fibers, in a tissue (38,39). We next determined the expression of fibrillar collagen family members by real-time PCR. The expression of collagen type I and type V did not change across groups (Fig. 3). The expressions of collagen III and IV in the uterus were significantly upregulated in the 0.15 and 1.5 ppm groups compared to the control, indicating higher levels of collagen expression in response to long-term phthalate exposure.

**Figure 3:**
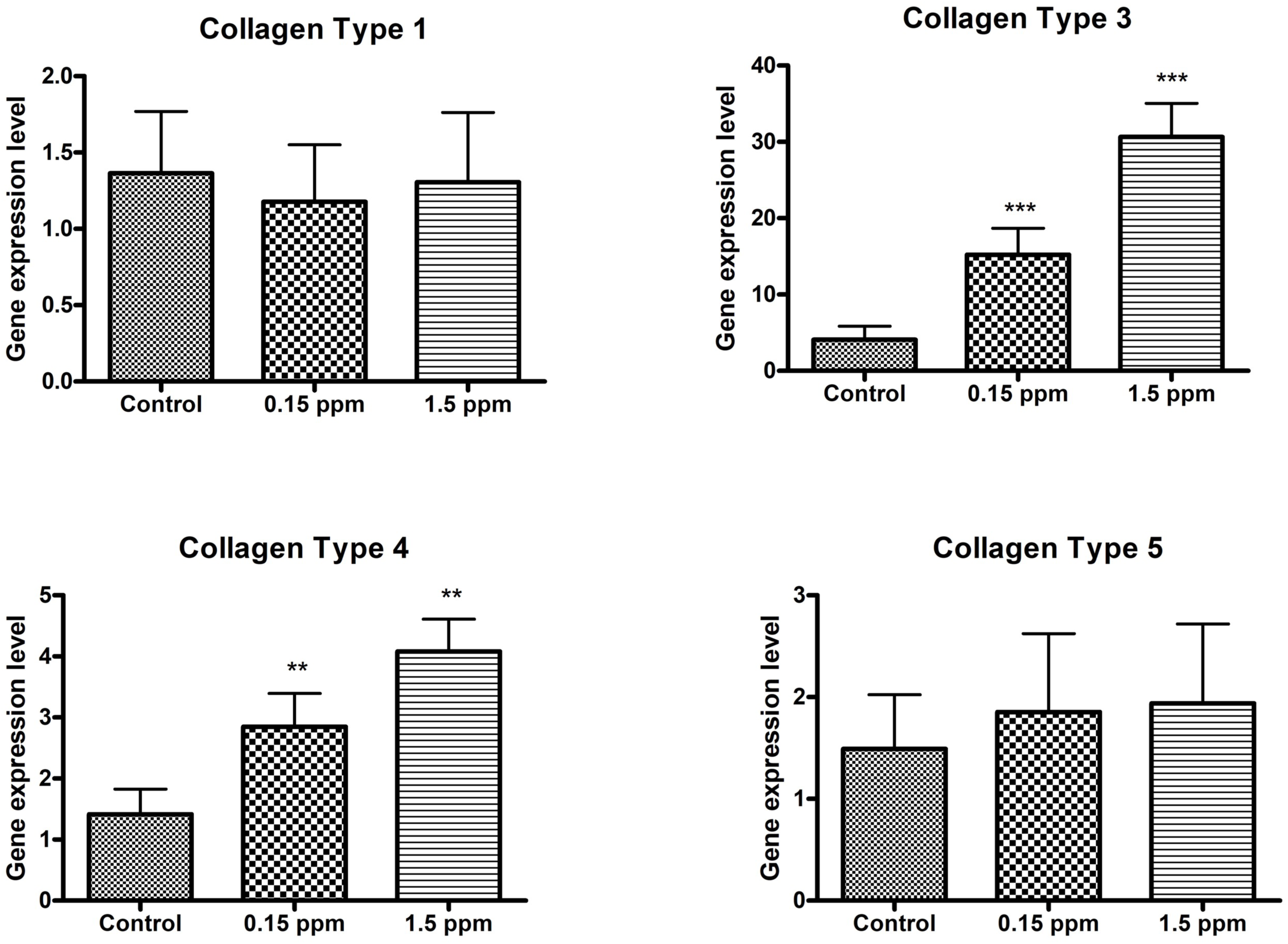
Expression of Collagen family members in the uterus. Uteri collected at 6 months were used for quantitative real-time polymerase chain reaction (qPCR) analysis. Uterine RNA from control groups, 0.15 ppm groups, and 1.5 ppm groups were subjected to qPCR using primers specific for *Collagen 1*, *Collagen 3*, *Collagen 4*, and *Collagen 5*. *36b4* was used as the internal control. Data represent mean ± SEM from six separate samples. Asterisks indicate statistically significant differences (\*\**P < 0*.01, \*\*\**P < 0*.001)

### 3.4. Elevated collagen deposition in the uterus upon exposure to a phthalate mixture

We then evaluated collagen deposition using Masson’s trichrome and Picrosirius red stain. In the Masson’s trichrome stain, blue color marks collagen. Our studies revealed more blue color, indicating elevated collagen in the phthalate-treated groups (Fig. 4, panels a-c). In the Picrosirius red stain, the collagen is marked by the red color and the intensity of red color indicates the amount of collagen content. We noted that in the control group, the collagen fibers were barely visible, but the phthalate-exposed samples exhibited pronounced red color indicating increased collagen deposition in the uterus (Fig. 4, panels d-f). Collectively, these results indicated an onset of uterine fibrosis upon long-term exposure to a mixture of phthalates.

**Figure 4:**
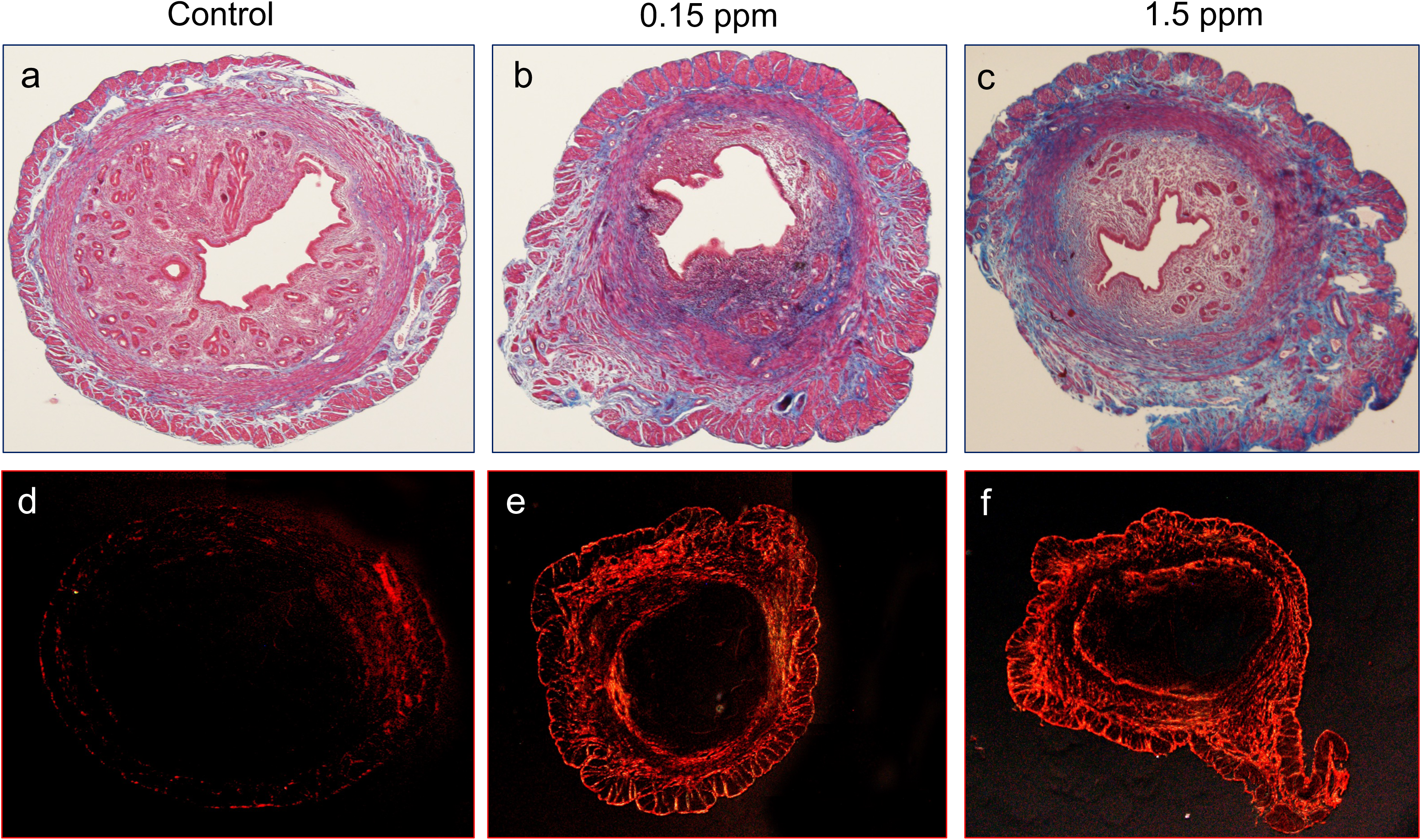
Exposure to a phthalate mixture leads to elevated deposition of collagen in the uterus. Upper: Mason trichrome staining of uterine cross sections of a control mouse (a), a 0.15 ppm group mouse (b), and a 1.5 ppm group mouse (c). Uterine samples were analyzed at 6 months. Collagen is marked by blue color. N = 3 per group. The representative images from the control and phthalate mixture treated groups are shown. Lower: Picro-Sirius red staining of uterine cross-sections of a control mouse (c), a 0.15 ppm group mouse group (d), and a 1.5 ppm group mouse (e). N = 3 per group. The representative images from the control and phthalate mixture treated groups are shown.

### 3.5. Exposure to a phthalate mixture leads to disorganized collagen in the uterus

Fibrotic states in a tissue are characterized by deposition of excessive collagen and disorganization of collagen fibers. Due to inflammation in fibrotic tissues, collagen fibers become disorganized and the new collagen fibers get deposited in a disordered manner, ultimately leading to increased stiffness of the tissue. We next performed an in-depth analysis of collagen fiber organization in phthalate mixture-exposed and unexposed uteri by performing Second Harmonic Generation (SHG) imaging. As shown in Fig. 5A, the collagen fibers in control samples were mostly parallel in their relative arrangement (panel b, control). In contrast, the collagen fibers were more disorganized in phthalate-treated uteri (panels b, treated).

**Figure 5:**
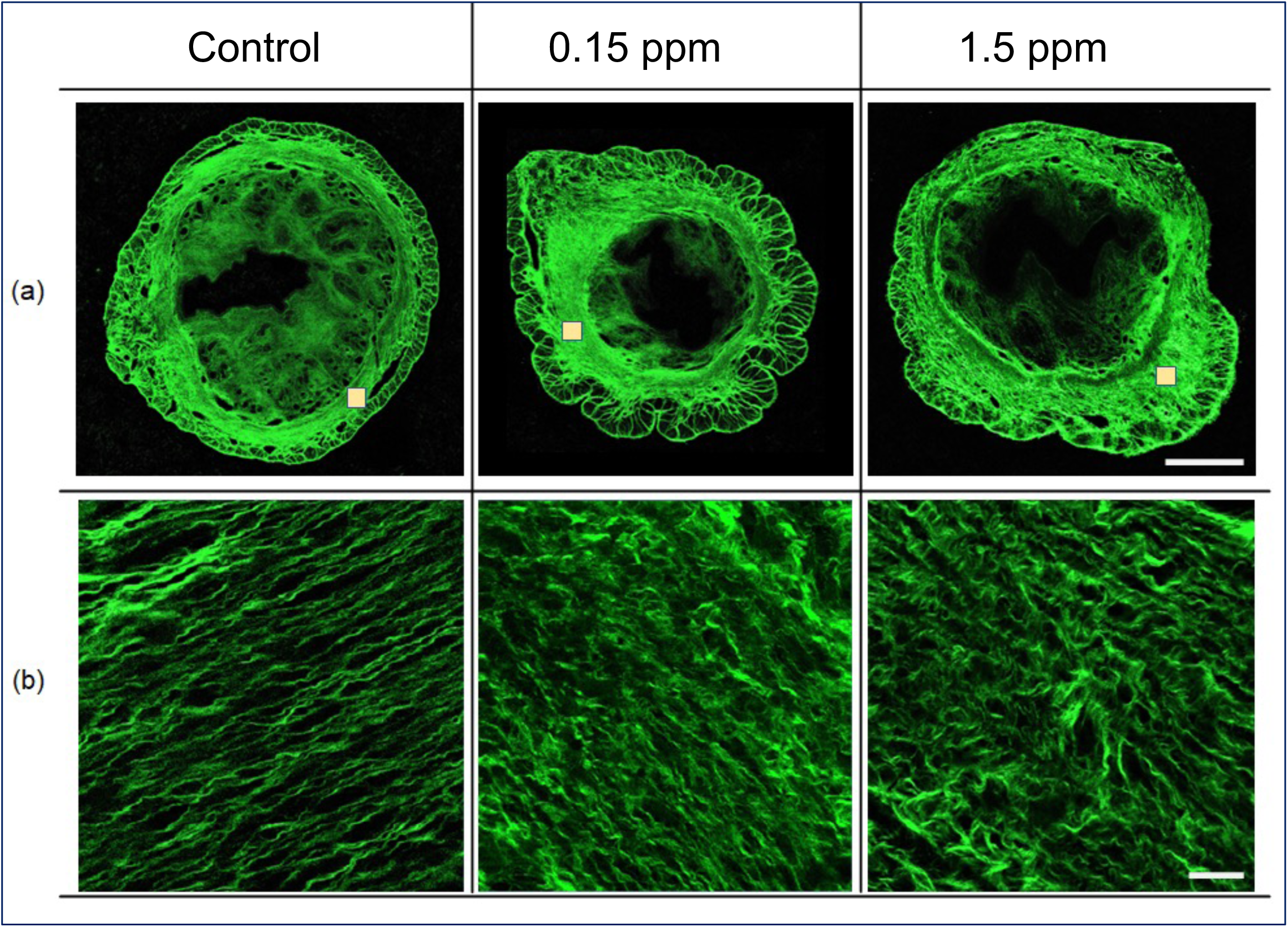
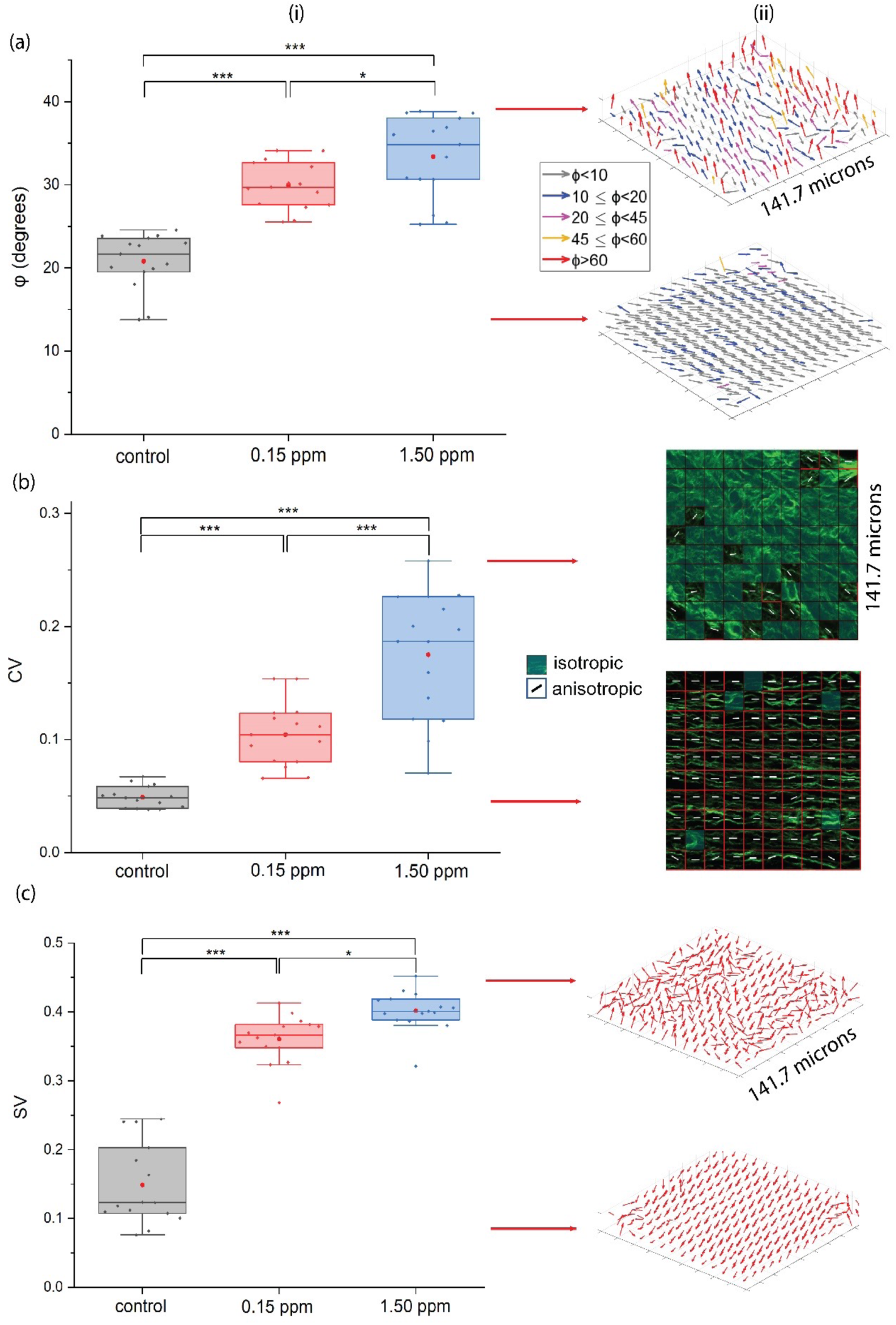
Exposure to a phthalate mixture leads to disorganized collagen in the uterus shown in mouse using Second Harmonic Generation (SHG) imaging. A (a): Uterine cross sections of a control mouse, a 0.15 ppm group mouse, and a 1.5 ppm group mouse. Scale bar is 500 microns. Uterine samples were analyzed at 6 months. Collagen is marked by green color. N = 3 per group. (b): High magnification images of the yellow boxes in the upper panels. Scale bar is 20 microns. Note significantly disorganized collagen fibers in phthalate-treated uteri. B. The extent of collagen fiber disorganization upon phthalate mixture exposure was determined quantitative analysis of SHG images. (a) Out-of-plane fiber angle is indicated by ϕ, (b) dispersion of the fiber orientations in the imaging plane is indicated by circular variance (CV), and (c) dispersion of fiber orientations in 3D is indicated by spherical variance (SV). N=3 per group. (i) Numerical value of SHG parameter for each group. (ii) Representation of collagen fibers showing vectors in 3D for (a) ϕ and (c) SV and in 2D for (b) CV. A one-way ANOVA followed by a post hoc Tukey analysis indicated significant differences. Three significance levels (p<0.05, p<0.01, p<0.001) were used for the analysis.

### 3.6. Quantitation of collagen fibers

We next quantitated the extent of collagen fiber disorganization in phthalate-exposed uteri by determining the SHG parameters including out-of-plane fiber angle (ϕ), circular variance (CV), and spherical variance (SV) (Fig. 5B). ϕ represents the average angle of the collagen fibers with respect to the x–y imaging plane, where 0° and 90° indicate in-plane and out-of-plane, respectively. CV is the dispersion of the fiber orientations in the imaging plane with a value between 0 (aligned fibers) and 1 (randomly oriented fibers). Similarly, SV describes the dispersion of fiber orientations in 3D ranging from 0 to 0.5. Fibers fully aligned in the imaging plane would have a low value for CV, a volumetric stack of imaging planes with fibers of similar alignment in all the imaging planes would have a low SV, and a stack with fibers oriented within the imaging planes of the volumetric stack would have a low ϕ. Our analysis of SHG these SHG parameters revealed significant disorganization of collagen fibers upon phthalate mixture treatment. Further, the 3D analysis of the fiber microstructure indicated that the treatment caused fibers to become significantly more out of plane, indicated by ϕ increase, and significantly more disorganized in 3D, determined by SV increase.

### 3.7. Measurement of tissue stiffness

Finally, we performed nanoindentation and co-registration of SHG images to identify the local mechanical properties of the tissue (Fig. 6). We found a significant increase in uterine stiffness when mice consumed 0.15 ppm and 1.5 ppm of phthalate mixture. There was also more variability in the indentation data with increasing consumption of phthalate mixture.

**Figure 6:**
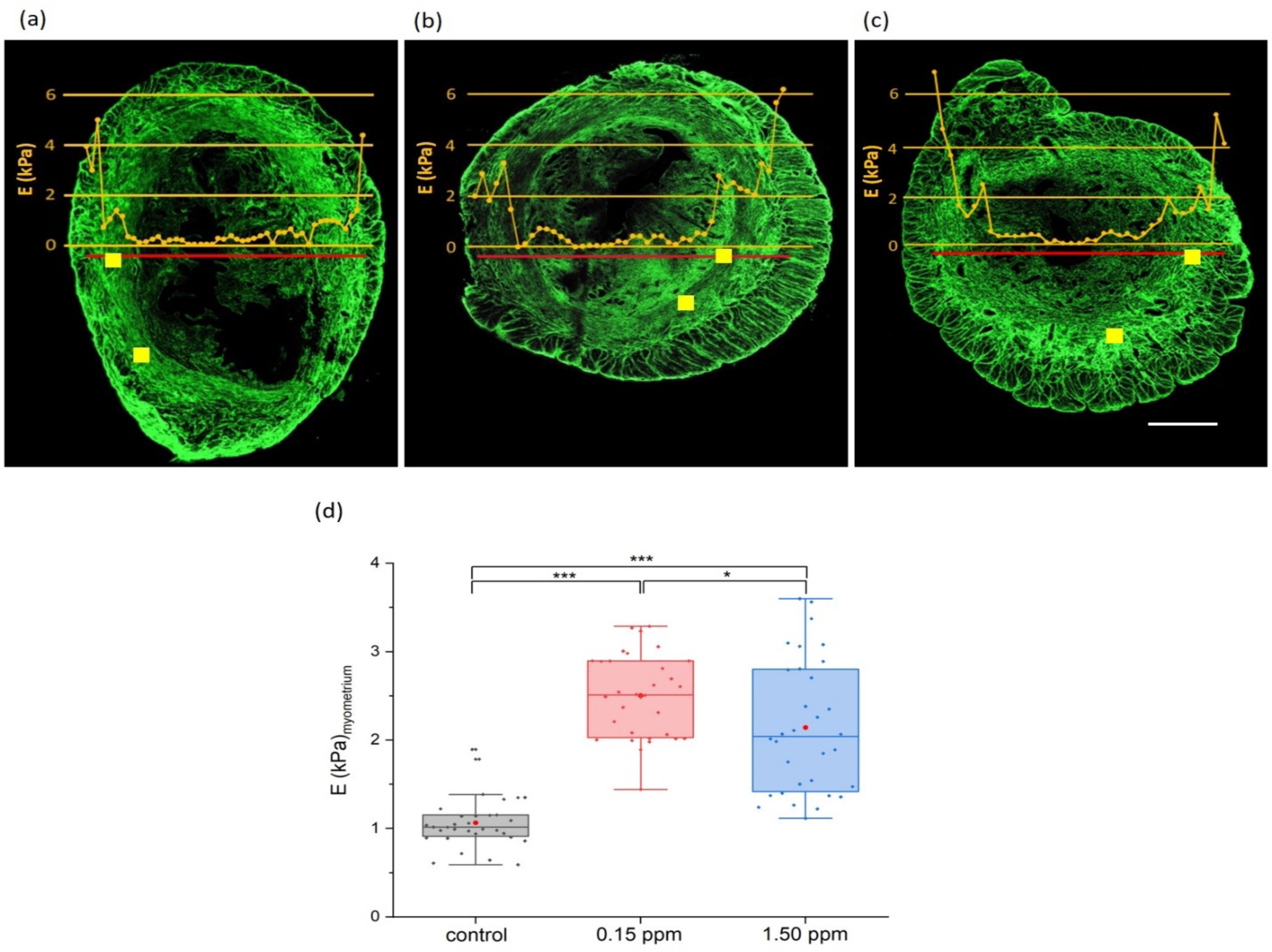
Exposure to a phthalate mixture leads to change in local uterine stiffness. The bright-field images from the nanoindention experiments and the SHG images were co-registered semi-manually using the tissue boundaries as fiducial references. The indentation data measuring modulus, a measure of stiffness, were overlaid on the SHG images using the coordinates taken from the two sets of bright-field images for the (a) control, (b) 0.15 ppm group mouse, and (c) 1.5 ppm group mouse. The marked yellow rectangular regions are representative regions chosen for measuring the stiffness of the myometrium. Data across groups compared in (d) and all groups are significantly different.

## 4. DISCUSSION

The aim of this study was to determine the effects of chronic exposure to a mixture of phthalates on the uterus. A previous study reported that a prenatal exposure to this phthalate mixture leads to multigenerational and transgenerational effects on uterine morphology in mice (40). However, human exposure to phthalates occurs daily over a lifetime. In this study, mice at six weeks of age were administered the phthalate mixture continuously for six months. Further, mice were exposed to phthalates via chow, which is relevant to human phthalate exposure. Collectively, our results show that long-term exposure to phthalate mixture via diet leads to fibrosis and local stiffening of the uterus.

A previous study showed that prenatal exposure to the phthalate mixture led to large cystic and dilated endometrial glands, higher incidence of myometrium intrusion into the endometrium, intrusion of endometrial tissues into the myometrium and myofibroblasts surrounding the glands (40). These results suggested an association between phthalate mixture exposure and adenomyosis or fibroids. Interestingly, we did not observe uterine adenomyosis or fibroids when mice consumed phthalate mixture for 6 months. However, we found an onset of uterine fibrosis in 6-month-old mice upon chronic exposure to phthalate mixture, indicating premature reproductive aging.

Tissue fibrosis, the excessive or dysregulated deposition of extracellular matrix (ECM) proteins, is a common pathophysiological condition of several diseases (38,39). TGF-β, a pleiotropic cytokine that plays an important role in regulating various cellular processes, is extensively implicated in fibrotic responses (38,39). The RNA-seq analysis revealed that the expression of three TGF-β isoforms (TGF-β1, - β2, and –β3), increase in the phthalate mixture exposed uteri. Consistent with the RNA-seq analysis, we observed significant increase in the level of TGF-β1 mRNAs in both 0.15 ppm and 1.5 ppm groups. TGF-β upon binding to the receptors TGF-β receptor 1 and 2 (TGFR1/2), phosphorylates Smad2 and Smad3, which complex with Smad4 and translocate to the nucleus. The Smad3 component of the complex binds directly to gene promoters to induce transcription of profibrotic molecules, including collagen (39,41). In addition, TGF-β orchestrates both pro- and anti-inflammatory responses in a cell and context-dependent manner. The RNA-seq analysis revealed that in the uterine context, NF-κB inflammatory pathway, which promotes the production of pro-inflammatory cytokines is activated upon exposure to the mixture of phthalates (39). However, the mechanism by which phthalates promote expression of genes in the TGF-β pathway requires further investigation.

We also observed a significant increase in the level of biglycan, a small leucine-rich proteoglycan that participates in the production of excess extracellular matrix and is related to fibrosis in many organs (42). Studies have shown that biglycan can enhance TGF-β signaling by increasing the binding of TGF-β to its receptors and promoting the formation of the TGF-β receptor complex. Additionally, biglycan can regulate TGF-β activity by modulating the expression and activity of TGF-β binding proteins. In addition to biglycan, integrin beta 1 can directly interact with the TGF-β receptor and activate downstream signaling pathways, including the Smad pathway. We also found an upregulation of Integrin beta 1 in response to phthalate mixture treatment compared to control. Interestingly, the phthalate-induced increases in biglycan and integrin beta 1 expression were observed in the lower 0.15 ppm group only. It is important to note that the relationship between the phthalate dose and effect occurs in a non-monotonic, dose-response-dependent manner. Despite the important implications that this phenomenon presents for the toxicological sciences, the molecular mechanisms underlying the non-monotonic dose-response for phthalates remain poorly understood.

Elevated expression of collagen mRNAs and deposition of collagen proteins indicated by Mason trichrome stain (blue color), Picrosirius red stain (red color), and SHG imaging indicate an onset of fibrosis in phthalate mixture treated uteri. Fibrosis is accompanied by inflammation, and it progresses due to interactions of inflammatory cytokines and fibrotic factors including collagen (39). It is interesting to note that pro-inflammatory NF-κB pathway is activated downstream of TGF-β signaling in the phthalate mixture treated uteri. Uterine fibrosis is an age-related change in rodents and our results indicate that exposure to phthalates causes premature reproductive aging. In women, uterine surgery or infections can cause fibrosis as a response to intrauterine adhesion (43). It is estimated that over 170 million women are affected by fibrosis, which may result in infertility and recurrent pregnancy loss (43).

Our studies reveal that phthalate mixture exposure led to excess collagen deposition and caused changes in myometrial collagen microstructural organization and stiffness. Exposure caused a significant change in orientation of collagen fibers from a uniform, circumferential pattern to a disorganized state. Notably, a uniform, circumferential pattern of collagen helps to provide support and elasticity to the uterus during pregnancy and disorganized collagen fibers can cause disruptions in these physiological processes. While little is known about the exact mechanisms that result in disorganized collagen in fibrotic state, there is increasing evidence that excessive deposition of collagen fibers in fibrosis can lead to increased stiffness and reduced elasticity of the tissue (44). A visibly higher density of collagen after phthalate mixture exposure likely causes the higher myometrial stiffness. The disorganized microstructure causing a higher stiffness may seem counterintuitive. However, the disorganization is partly caused by the out-of-plane fibers, which are approximately parallel to the indentation axis when indenting onto the transverse cross-section. Our previous work showed that fibrous samples are stiffer when indented parallel to the collagen fibers (45).

In summary, this study to our knowledge is the first to investigate the long-term effects of phthalate exposure on uterine homeostasis and function. Our results showed that chronic exposure to a phthalate mixture leads to an upregulation of TGF-β signaling and a premature onset of fibrosis in the uterus. Additionally, using nanoindentation and SHG imaging, our data demonstrate that phthalate mixture exposure increases local uterine stiffness due to the excess collagen. Future studies would examine the mechanisms by which chronic exposure to phthalates alters the expression of genes involved in the TGF-β pathway and disrupts uterine homeostasis.

## Acknowledgements

This work was supported by NIH (R01 ES032163, R01 ES034112, and T32 ES007326).

